# Converting Organic Municipal Solid Waste into Volatile Fatty Acids and Biogas: Experimental Pilot and Batch Studies with Statistical Analysis

**DOI:** 10.1101/2023.06.21.545938

**Authors:** Hojjat Borhany

## Abstract

**Background:** Italy can augment its profit from biorefinery products by altering the operation of digesters or different designs to obtain more precious bio-products like volatile fatty acids (VFAs) than biogas from organic municipal solid waste. In this context, recognizing the process stability and outputs through operational interventions and its technical and economic feasibility is a critical issue. Hence, the present study is an anaerobic digester in Treviso in northern Italy.

**Objectives:** This research compares the novel line, consisting of pretreatment, acidogenic fermentation, and anaerobic digestion, with single-step anaerobic digestion regarding financial profit and surplus energy. Therefore, a mass flow model was created and refined based on the outputs from the experimental and numerical studies. These studies examine the influence of hydraulic retention time (HRT), pretreatment, biochar addition, and fine-tuned feedstock/inoculum (FS/IN) ratio on bio-products and operational parameters.

**Methods:** VFA concentration, VFA weight ratio distribution (VWRD), and biogas yield were quantified by gas chromatography. Then, a *t* test was conducted to analyze the significance of dissimilar HRTs in changing the VFA content. Further, a feasible biochar dosage was identified for an assumed FS/IN ratio with an adequate long HRT using the First-Order (FO) rate model. Accordingly, the parameters for a mass flow model were adopted for 70,000 population equivalents to determine the payback period and surplus energy for two scenarios. We also explored the effectiveness of amendments in improving the process kinetics.

**Results:** Both HRTs were identical concerning the VFA/SCOD (0.88 kg/kg) and VWRD: mainly acetic acid (40%), butyric acid (24%), and caproic acid (17%). However, a significantly higher mean VFA content was confirmed for HRT 4.5d than the quantity for HRT 3d (30.77 vs 27.66, g-SCOD/L) using a *t* test (t =- 2.68, df=8, ***P***=.03, CI=95%). In this research, 83% of the fermented volatile solids were converted into biogas to obtain a specific methane production of 0.133 (/kg−VS). While biochar addition improved only the maximum methane content by 20% (86% v/v), the feedstock/inoculum ratio of 0.3 (volatile solid basis) with thermal plus fermentative pretreatment improved the hydrolysis rate substantially (0.57 vs 0.07, 1/d). Furthermore, the biochar dosage of 0.12 (g-biochar/g-VS) with HRT 20d was identified as a feasible solution. Principally, our novel line payback period would be almost 2 years with surplus energy of +2251 (MJ/d) compared to 45 years and +21567 (MJ/d) for single-step anaerobic digestion.

**Conclusions:** This research elaborated on the advantage of the refined novel line over the single-step anaerobic digestion and confirmed its financial and technical feasibility. Further, changing HRT and other amendments significantly raised the VFA concentration and the process kinetics and stability.

## Introduction

The European Union (EU) annually generated about 110 million tons of organic waste in 2006, which excluded slurry and manure. This waste mainly came from the food industry (33%), agriculture and hunting (30%), and households (20%) [1]. Current Italian legislation forbids landfilling organic waste but requires treating it through biological and thermal processes like anaerobic digestion, composting, and incineration with high disposal costs for secondary waste flux (75-125 €/ton) [2]. Under the pressure of exhaustible natural exploitation and increasing organic waste, the European Commission approved the circular economy action plan to promote sustainable recovery methods to reduce the secondary waste flux. The techniques recommended in the circular economy context assume a “take-use-reuse” viewpoint. Such an approach wants to close the circuit of cycles, extend product life, and treat the wastes as precious recyclable materials [3,4]. In this respect, the EU states have deployed biological processes such as anaerobic digestion to gain either platform chemicals like volatile fatty acids (VFAs) or biogas from organic wastes produced in urban areas [5–9]. These products are extremely valuable in the era of environmental disasters with several consequences (e.g., climate change) since they are renewable, sustainable, carbon-neutral, and compatible with current fossil-based fuel infrastructures [10].

The recent studies aimed at finding a sequential reclaiming route to obtain various bioproducts such as VFAs and biohydrogen with a higher added-value market than bio-methane at distinct steps to either redesign the existing plants or integrate them into biorefinery platforms [11,12]. Various biological processes can convert different feedstock (e.g., edible sugary crops, oil-bearing crops, livestock, waste sludge (WS), and food waste) into a range of biofuels, including bioethanol, biodiesel, biomethane, and biohydrogen [10,13,14]. Biofuel production from edible crops is quite controversial in terms of food supply, ethical quandary, and insecure supply chain. However, food waste, WS, and livestock are omnipresent in urban and rural areas without widespread deployment in a biorefinery scheme. Accordingly, this research aims to convert organic municipal solid waste (OMSW), mainly from food waste, into VFAs and biogas.

This study examines the biological recovery route for OMSW for the potential beneficial bioproducts and technical feasibility. This effort includes three steps: pretreatment, mesophilic acidogenic fermentation, and anaerobic digestion. Specifically, it is endeavored to conceive how to make the process more profitable and practicable through operational amendments that change the share of methanogenesis and acidogenic routes in the final products (VFAs and Biogas) [9] and lower the costs of the process in terms of energy and water consumption. Hence, determining a reasonably priced process with a desirable VFA-rich stream from acidogenic fermentation and a high methane yield from methanogenesis [15] could ultimately encourage full-scale commercialization. VFAs typically serve as platform chemicals for many processes (e.g., biopolymer synthesis of polyhydroxyalkanoates (PHAs) [16–19]), which could be later recovered through biological processes to close the material life cycle.

The major bottleneck in anaerobic digestion of biowaste is at the hydrolysis step. Such a problem could be relieved by various methods such as pretreatment, optimized feedstock/inoculum (FS/IN) ratio, and carbonaceous material addition, including biochar [20–22]. The latter method was recently realized to have numerous benefits to the process, such as improving the process stability, accelerating the process rate, buffering potency and alkalinity, inhibitors adsorption, enriched microbial functionality, and electron transfer mechanism. As a result, it could improve CH_4_ generation by fostering hydrolysis, acetogenesis, and methanogenesis [23]. The residual solids out of the multistep line of pretreatment followed by acidogenic fermentation plus anaerobic digestion can be used in a pyrolysis line for biochar and biofuel production to further lower the secondary waste flux [24]. This strategy provides several benefits, such as combating climate change and global soil degradation and addressing the rising energy demand.

This study compares the multistep route of pretreatment, acidogenic fermentation, and anaerobic digestion with the existing method of single-step anaerobic digestion for valorizing OMSW in the Treviso wastewater treatment plant (WWTP) in terms of financial profit and technical feasibility. In this context, the present research has the ultimate goals of facilitating the entrance of the process into the market and further closure of the cycle of organic material. Accordingly, it assesses several suggestions, such as hydraulic retention time (HRT) variation, pretreatment, biochar addition, and adjusted FS/IN ratio to enhance the bio-products and decline the involved costs. To this end, their effects on the process were quantified through experimental tests, confirming their significance through statistical analysis. Later, the payback period, amount of surplus energy, and volatile solids (VS) destruction for the mentioned scenarios were determined using a mass balance model refined according to the laboratory studies. The boundary condition parameters for energy conversion and costs were assumed according to previous studies and experts’ knowledge, respectively. To the best of the author’s knowledge, this paper is novel in presenting a robust framework to assess a groundbreaking proposition for the valorization of OMSW financially and technically. Overall, it is concluded that our line is viable technically and overtakes the conventional methods financially.

## Methods

### Biorefinery process scheme and experimental studies

(Figure 1) presents the hypothesized biorefinery process line in this research. It comprises screw-pressing, pretreatment unit, mesophilic acidogenic fermentation, solid-liquid separator, and mesophilic anaerobic digestion. The two sectors of biopolymer production and pyrolysis were exhibited differently since no mass and energy flow was considered for them, and only the possible end goals for the secondary stream were shown.

**Figure 1.**
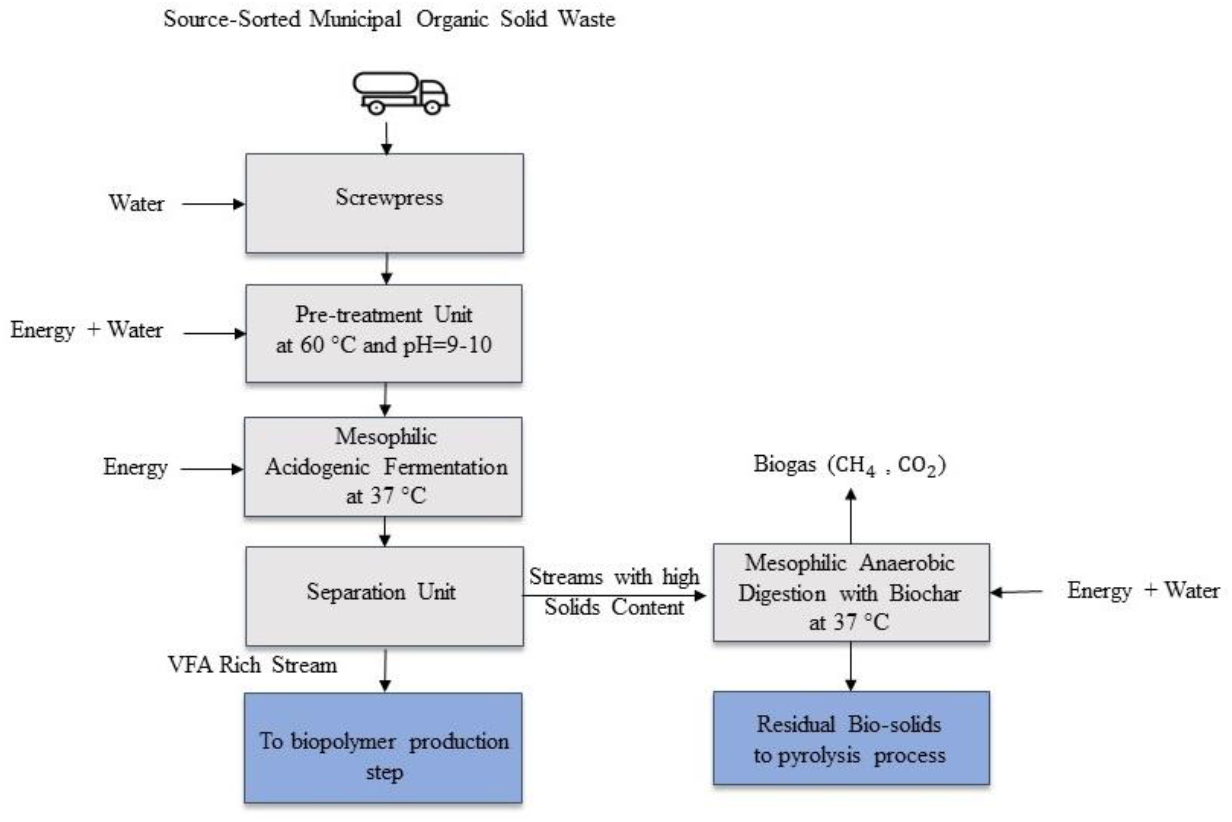
Schematic of the multistep of pretreatment, acidogenic fermentation, and anaerobic digestion for VFAs and biogas production from the OMSW

After and before the pretreatment, the feedstock for different parameters was characterized from time to time. These parameters include the total solids (TS), VS, chemical oxygen demand (COD), soluble COD (SCOD), Total Kjeldahl Nitrogen (TKN), total phosphorous (P),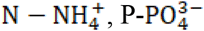, and VFA.

The arrived feedstock in the WWTP had already been mixed with the acidogenic fermentative inoculum, which initiated solubilizing and converting the organic solid matters into SCOD and VFAs in the transporter. Then, in the pretreatment unit, NaOH solution (40% kg/kg) was added to bring the pH to 9-10 and heated to 60° for 24 hours. Subsequently, the bio-mixture was fed manually into 5 L (operational volume of 4.5 L) continuously stirred pilot acidogenic fermenter operated at the given conditions (Table 1). Its high alkalinity maintained pH during the acidogenic fermentation in the optimal range. Further, the mixture was blended mechanically, and the whole system was kept in the oven to hold the temperature constant at 37°. The output was sampled frequently during the week, and the samples were centrifuged to obtain the supernatant to measure pH, SCOD, VFA,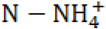, and 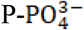 A tiny fraction of the residual solid part was used to characterize solids like COD, P, and TKN, and the rest was kept in the freezer to apply the biomethane potential test (BMP).

**Table 1.** The operational parameters of the mesophilic acidogenic fermenter.

| HRT (Days) | OLR (kg-VS/m<sup>3</sup> . d) | Temperature (°C) | pH |
| --- | --- | --- | --- |
| 4.5 | 6.89 | 37 | 6.13-6.96 |
| 3 | 10.33 | 37 | 5.95-7.55 |

The VS and TS characterization were performed in 105° and 550° ovens for 24 hours. Except for VFA, all the remaining analyses (including COD measurements) followed the standard methods for examining water and wastewater [25]. The method described in the A and D sections No. 5220 for COD quantification was employed. These methods are named “Closed Reflux, Titrimetric Method” and “Closed Reflux, Colorimetric Method” for the solid and liquid phases, respectively [25]. For the liquid, the samples were filtered after being centrifuged at 4500 rpm for 5 minutes, and before the analysis, the supernatant was filtered with a 0.45 (μm) cellulose filter (Whatman). For the solid, acidic digestion was performed at 220° and a high pressure of 2 atmospheres to destroy the 0.2 g of solid matrix for 2 hours. Afterward, the COD was measured in the solution using titration by ferrous ammonium sulfate as described in the standard methods. Our limit of detection (LOD) was 50-500 (mg-COD/L) for the calorimetric method and 40-400 (mg-COD/L) for titrimetric method. In this research, dilution was done for a high concentration value, which is beyond the considered LOD.

The VFA was quantified using a gas chromatograph (Agilent Technology TM, 6890N) equipped with a flame ionization detector (with a temperature of 100°), a fused silica capillary column, DB-FFAP (15 m × 0.53 mm × 0.5 mm thickness of the film), and hydrogen as the gas carrier. The chromatographic run was performed by increasing the temperature from 80 to 200° at a rate of 10 °/min. The VFA samples were centrifuged at 4500 rpm for 5 minutes, and before the analysis, the supernatant was filtered with a 0.22-μm cellulose filter (Whatman).

In the BMP, the effect of biochar addition was observed for 3 diverse dosages (0-0.12-0.24, g-biochar/g-VS) on the biomethane volume, content, and production kinetics in the mesophilic condition using four sets of BMP. The anaerobic condition was assured in bottles just by sealing them after filling without any flushing with nitrogen or,co_2_ since we had known that the oxygen transfer at the surface of waste stream was impossible as it contained a high TS and SCOD. This type of procedure was adopted in our lab and has been conducted for years. The tests were conducted with a total number of 8 bottles of 250 mL (working volume of 215 mL). The biochar was synthesized by a local supplier, and its main physical and chemical features are reported in the supplementary document (Table 1). It was ground into microparticles and kept under the dried condition at room temperature before being added to the bottles. Further, the inoculum for the BMP was collected from 2,300 completely stirred anaerobic digester treating thickened waste sludge (WS) and squeezed organic municipal solid waste mixture under the mesophilic condition at an OLR of 1.8-2.0 (kg-VS/) in the treatment plant. The inoculum was added to the feedstock (residual solid from acidogenic fermentation) based on the weight ratio of 0.3 (FS g-VS/IN g-VS). The TS and VS contents in the bottles (i.e., inoculum and feedstock) were 133 (g/kg) and 17.6 (g/kg), respectively.

The experiments were conducted for each condition, namely only inoculum and either with or without biochar, in 2 bottles. The test was terminated after 25 days when the cumulative biogas production reached almost 89% of the final projected value. The biogas content was characterized by gas chromatography (for days 1, 4, 6, 10, 14, 16, 18, 21, and 25). Also, the values for the remaining days were filled through imputation using the K-Nearest Neighbors Algorithm (number of neighbors = 4 and weights = distance) [26]. The imputation code was provided in the supplementary documents. Then, the biogas and biomethane volumes were subtracted from the only inoculum to correct for the endogenous methane production, and both values were averaged for 2 bottles. Gas chromatography was performed using (Agilent Technology, TM 6890N) with an HP-PLOT MOLESIEVE column (30 m length, 0.53 mm ID × 25 mm film thickness) and a thermal conductivity detector (TCD) with argon as a carrier (79 ml/min).H_2_,CH_4_,O_2_, and N_2_were analyzed using a TCD at 250°. The inlet temperature was 120°, with constant pressure in the injection port (i.e., 70 kPa). Samples were taken using a gas-type syringe (200 µL). Once the entire sample was vaporized, peak separation occurred within the column at a constant temperature of 40° for 8 minutes. We didn’t plan on monitoring pH and other parameters like alkalinity, VFA, ammonia, and phosphate because the pH drop risk was negligible, and the biochar addition could provide a buffer capacity and adsorption of inhibitory compounds in the solution [27]. Moreover, a considerable part of the readily biodegradable COD of the feedstock was already converted to VFAs in the previous step. As a result, the process was easily controlled even in the transient condition when the risk of methanogenic inhibition was high [28].

### Statistical analysis and performance indicators

The performance indicators, including COD solubilization, VFA yield, ammonia and phosphate release, and VFA/SCOD ratio, were determined. These indicators characterize the mesophilic acidogenic fermentation on the days when the data were available, and the process reached the pseudo-steady state condition. In addition, the VFA weight ratio distribution (VWRD) was determined from the total VFA weight on the same day. The process stability was evaluated based on variations in daily VFA concentrations. The indicators were calculated, and the data were plotted using a Microsoft EXCEL spreadsheet. The formula for the performance parameters was reported in the supplementary documents. The exploratory data analysis and *t* test on VFA data were performed for the VFA concentration, yield, and VFA/SCOD ratio for two HRTs by the open-source program R (The R Foundation for Statistical Computing, version 3.5.0). We assumed that the 2 data sets are paired and have a normal distribution. The code is provided in the supplementary documents. The values for two HRTs to increase the VFA concentration in the outlet were selected based on our experience and process knowledge. According to this information, exceeding the HRT value of more than 3-5 days can bring the process into an anaerobic digestion step. As a result, the VFAs with high-added value markets are converted to biogas. Hence, the two HRTs of 3d and 4.5d were tried in pilot test, knowing that the VFA concentration would either increase or decrease linearly in this local region of operation.

For the BMP, two kinetic models were calibrated, namely the First-Order (FO) rate and Modified Gompertz (MG), to the biogas’ cumulative yield. Also, the specific methane/biogas production (SMP, SGP) plus maximum volumetric methane content (v/v %) were determined. Comparing these results could reveal how the biochar addition, FS/IN ratio of 0.3, and pretreatment improved the process in terms of rate and fostered methanogenesis. Such improvement is manifested through a higher hydrolysis rate, a shorter lag phase, and a higher maximum volumetric methane content. Besides, the biogas yield was determined in terms of (g-biogas/g-VS).

### Technical and economic assessment

This research sets up a mass flow analysis with parameters adopted for a municipality with 70,000 population equivalents (PE) for the two scenarios: (1) a line with pretreatment, mesophilic acidogenic fermentation followed by mesophilic anaerobic digestion and (2) a single-stage mesophilic anaerobic digestion as currently deployed in Treviso WWTP. This study focuses on water and energy preservation and increase the profit from VFA production in our conversion line through several refinements. They were tied with the HRT identified in the previous step, integration of our process knowledge of using the fine-tuned FS/IN ratio, and biochar addition in anaerobic digestion. Detailed information and calculations regarding the mass flow analysis are available in the supplementary documents in the EXCEL spreadsheet named “Mass Balance”. In the following, the full description of the two scenarios was expressed:

The two scenarios shared the first part of the model where the separated OMSW by a door-to-door collection system was screw-pressed and diluted with water to reach the TS of 280 (g/kg). Then, in the first scenario, adding sodium hydroxide solution (40% kg/kg) elevated the feedstock pH to 9-10. Afterward, the solution was heated at 60° for 24 hours in the pretreatment unit. Next, it was diluted and heated further before feeding into the mesophilic acidogenic fermenter based on the desirable HRT. The last part of the first scenario was the optimized anaerobic digestion of residual fermented solids. Specifically, the stability endowment by adding biochar to the anaerobic digestion could ultimately smooth running the process in a high OLR (low water dilution). Furthermore, FS/IN of 0.3 was applied to increase the kinetics rate with the pros of a decrease in digester volume, energy consumption, and capital cost. This finding is of significant importance in plants and zones with limited area, water, and energy.

In the second scenario, the screw-pressed feedstock was diluted and immediately fed into a mesophilic anaerobic digester for only biogas production.

It was assumed that the reactors transfer heat from the walls with the atmosphere and earth. Further, the biogas would be consumed in the combined heat and power units for electricity production with an overall efficiency of 0.4. In this research, the mass of VFAs and the net amount of energy production were accounted for as the source of income. Meanwhile, the corresponding costs were the operational expenditure, the mass of the water process, and the final residual solids to dispose of. Reference parameters for energy analysis and boundary conditions are given in (Table 2). The price of electricity was assumed to be 130 €/MWh. These two scenarios were compared to identify the most favorable one regarding surplus thermal energy and electricity or the shorter payback period.

**Table 2.**
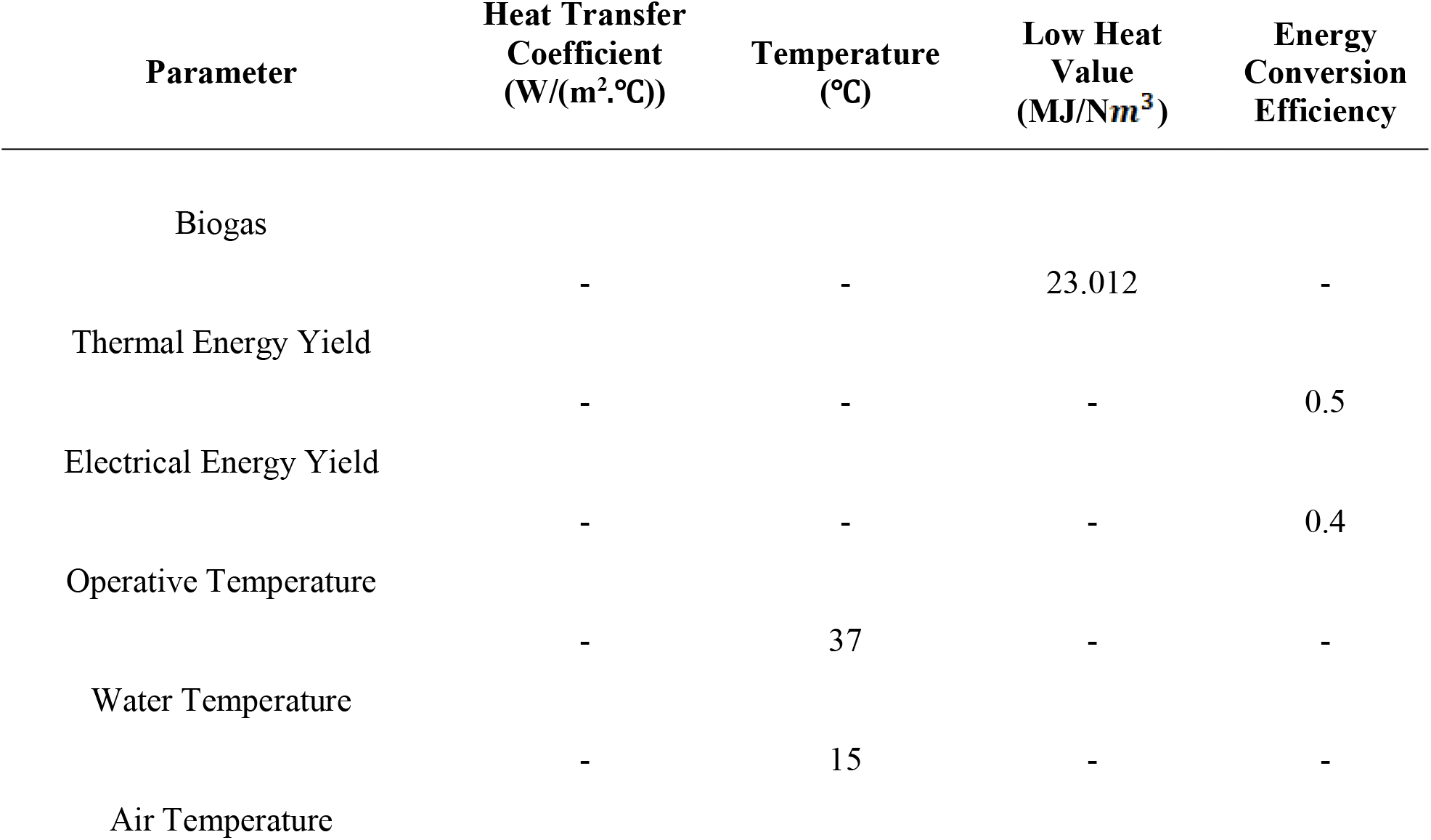
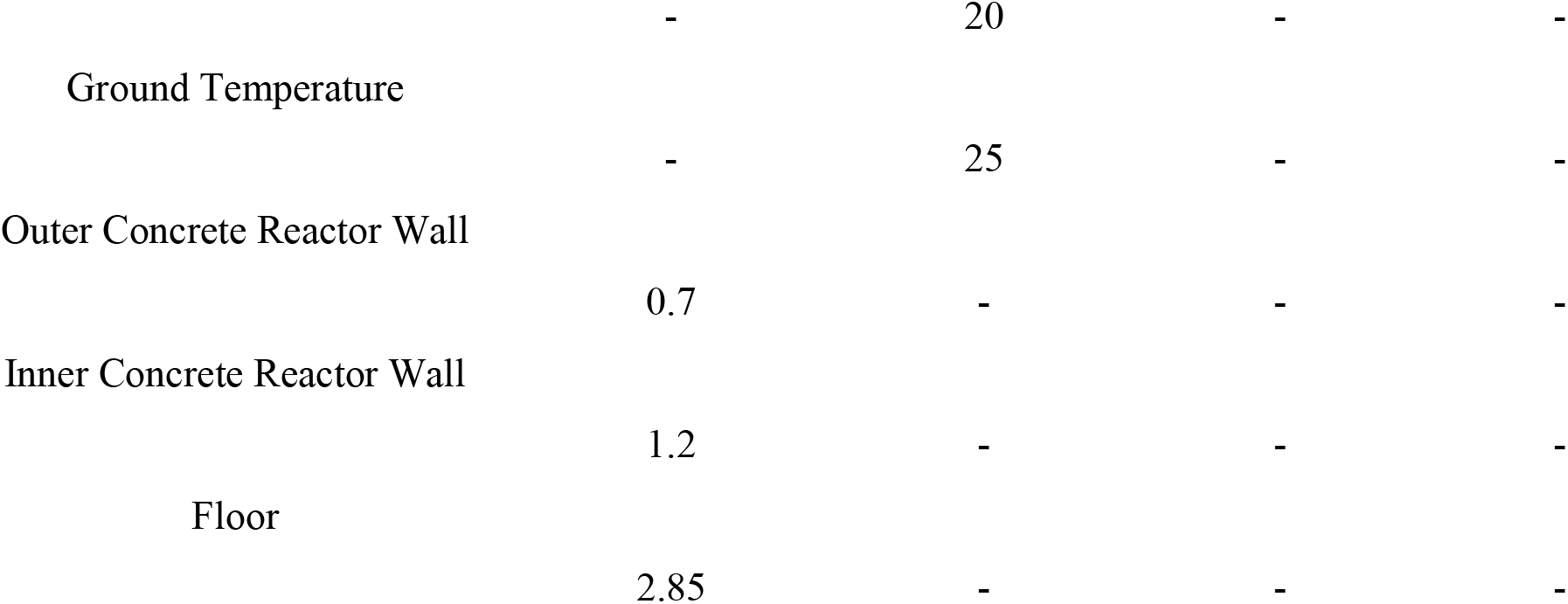
Reference parameters and boundary conditions for energy flow analysis.

## Results

### Biorefinery process scheme and experimental studies Composition and characteristics of the pretreated feedstock

The pretreated feedstock’s main physical and chemical characteristics were quite stable throughout the experiment (Table 3). The feedstock had an average TS content of 45 (g/kg) and VS content of 32 (g/kg). These values suggest that the biodegradable solids constituted 72% part of the TS, which could support the fermentation process. The chemical composition of the solid part was 12.9(g-N/kg-TS), 4 (g-P/kg-TS), and 565 (g-COD/kg-TS), which was in the range of the values reported for the typical OMSW in Italy [29]. The chemical composition of the liquid was 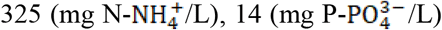, and 25.8 (g-SCOD/L). Further, the feedstock COD:N:P ratio was determined as 100/2.2/0.7, meaning that nutrients such as phosphor and nitrogen would be the limiting substrates in acidogenic fermentation [30]. In this regard, the slight level of VFA concentration at the level of 3.5 (g-SCOD/L) was due to acidogenic fermentation, which had been happening during transportation.

**Table 3.** Main physical-chemical features of the pretreated feedstock.

| Parameter | Weight Ratio<br>(g/kg) ± SD | Percentage (%)<br>± SD | Concentration<br>(mg/L) |
| --- | --- | --- | --- |
| TS<sup>a</sup> | 45±3.21 | - | - |
| VS<sup>a</sup> | 32±3.28 | - | - |
| TKN<sup>b</sup> | 12.9 | - | - |
| P<sup>b</sup> | 4 | - | - |
| COD<sup>b</sup> | 565 | - | - |
| COD: N: P | - | 100:2.2:0.7 | - |
| SCOD | - | - | 25,814 |
| N-$NH_4^+$ | - | - | 325 |
| P-$PO_4^{3-}$ | - | - | 14 |
| VFA<sup>b</sup> | - | - | 3,500 |
| VS/TS<sup>a</sup> | - | 72%±5% | - |

a: 3 measurements, b: measurements done for nitrogen, phosphor, and SCOD equivalents for TKN, P, COD, and VFA, respectively

### Statistical analysis and performance indicators Acidogenic fermentation

(Table 4) presents the main physical and chemical characteristics of the effluent and solid cake from the acidogenic fermenters. According to (Figure 2), the process reached a steady condition after 14 days, which was roughly 3 times HRTs (4.5 days). Both HRTs were similarly stable in terms of VFA concentration variation because of a negligible difference between standard deviations: 2.82 (g-SCOD/L) vs. 2.45 (g-SCOD/L). This value was almost 1% of the total VFA, and the VFA production continued for more than 3 weeks without any considerable issues. The lack of any change in this process is attributed to the initial high pH of 9-10, which supported the process by keeping the pH variation in the optimal range of 6-7.5 [31].

**Table 4.** Main physical-chemical features of the effluent and solid cake from mesophilic acidogenic fermentation.

| HRT (Days) | TS (g/kg)±SD | VS (g/kg)<br>±SD | VFA (g-SCOD/L)<br>±SD | pH±SD |
| --- | --- | --- | --- | --- |
| 4.5<sup>a</sup> | 43±5.15 | 23.6±2.07 | 30.77±2.82 | 6.56±0.25 |
| 3<sup>b</sup> | 38±4.55 | 25.8±1.5 | 27.67±2.45 | 6.7±0.45 |

a: 5 measurements for TS and VS, 9 measurements for VFA, 13 measurements for pH, b: 4 measurements for TS and VS, and 9 measurements for VFA, 13 measurements for pH

**Figure 2.**
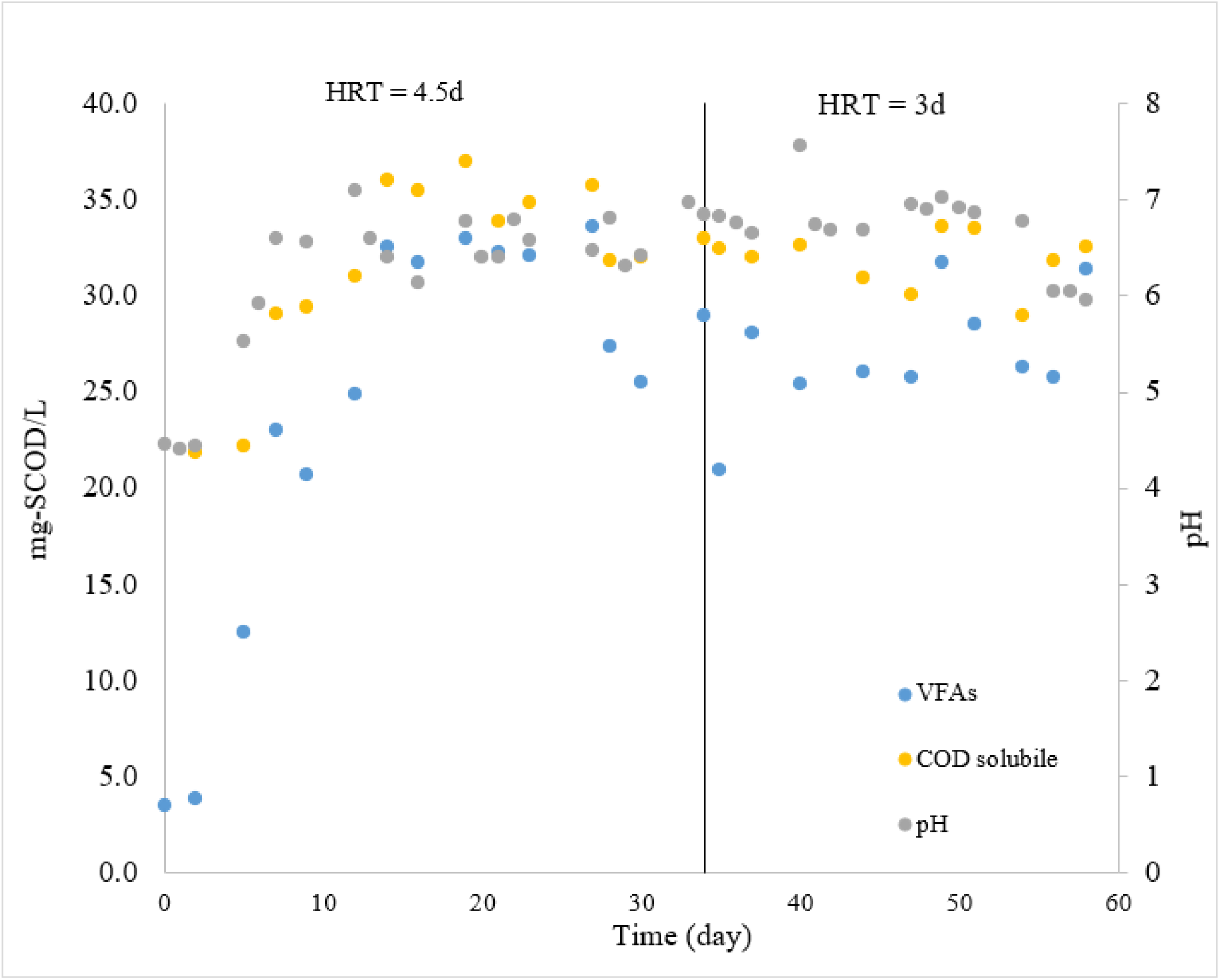
VFA, SCOD, and pH for mesophilic acidogenic fermentation

Based on the *t* test results (t=-2.68, df=8, ***P***=.03, CI=95%), it is verified that the mean VFA concentration for HRT 4.5d was significantly higher than the value for 3d (30.77 vs 27.67, g-SCOD/L). A similar statistical analysis (t=-0.99, df=8, ***P***=.35, CI=95%) for the VFA/SCOD ratio rejected the significance of a higher mean value of 0.892 for HRT 4.5d than 3d with the value of 0.87. The possible range of values for the VFA concentrations and VFA/SCOD, which cover 99% and 50% of the data for the 2 HRTs, were depicted by the boxplots in (Figures 3) and (Figure 4), respectively.

**Figure 3.**
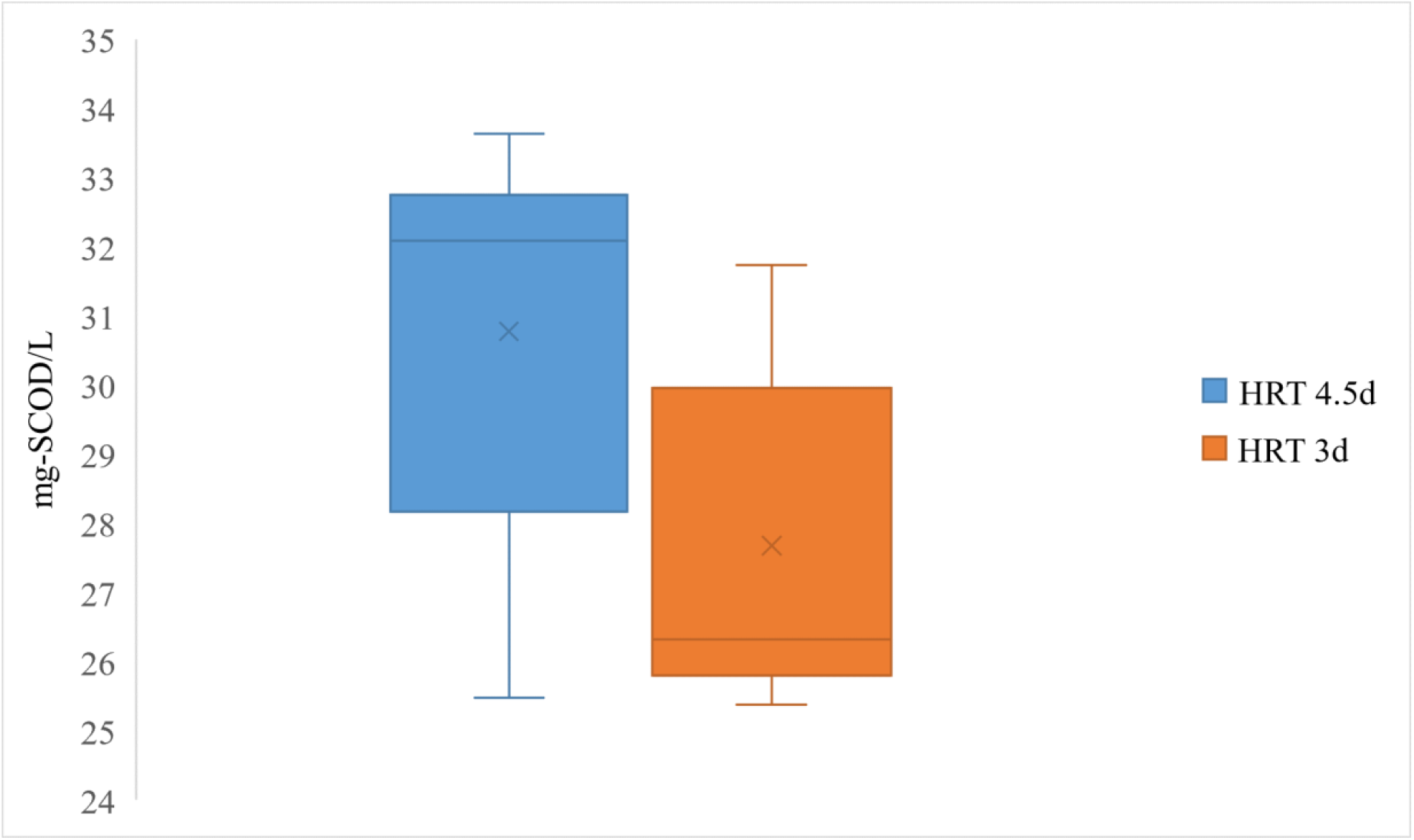
Boxplot of VFA concentrations for mesophilic acidogenic fermentation

**Figure 4.**
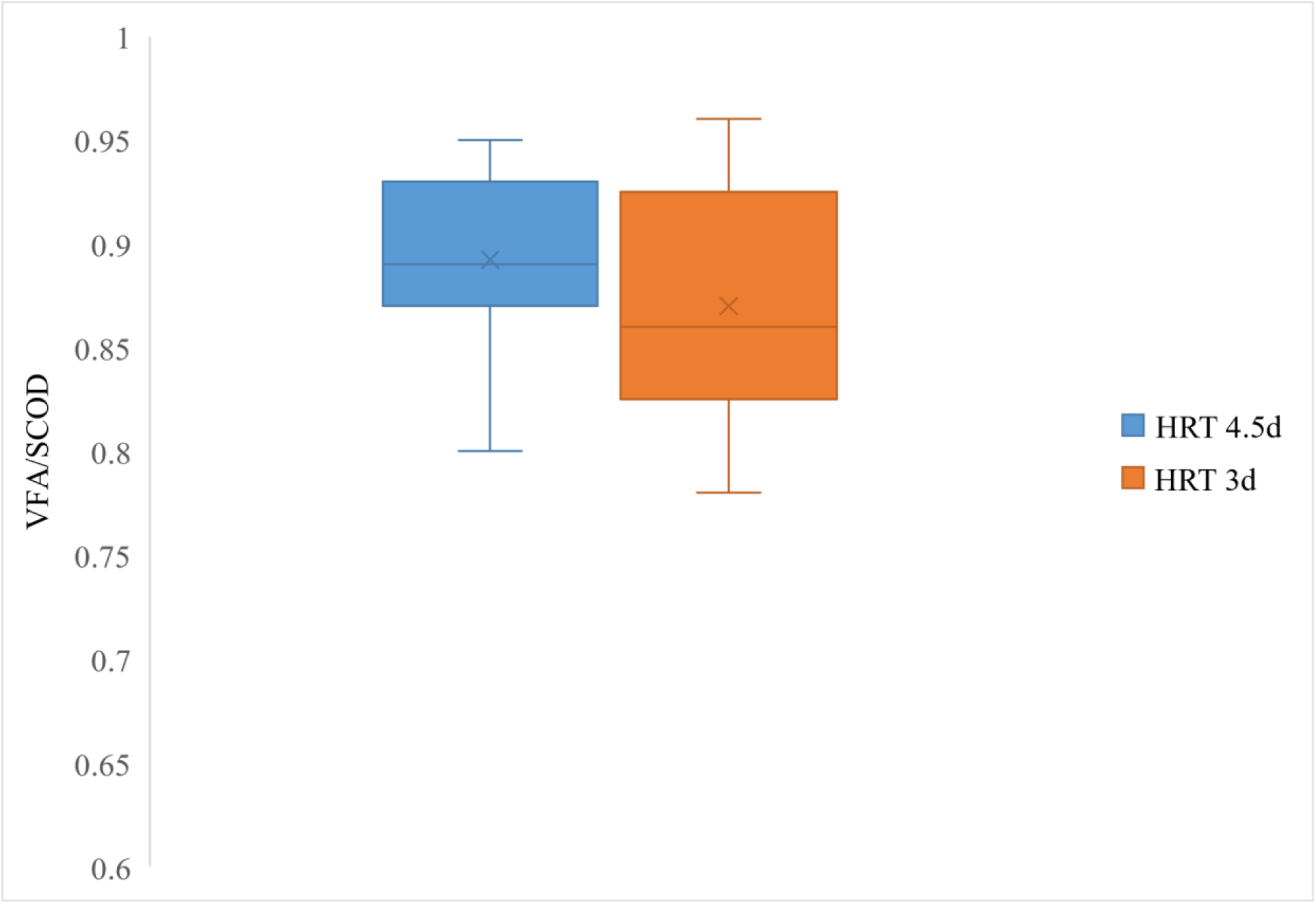
Boxplot of VFA/SCOD ratios for mesophilic acidogenic fermentation

Performance parameters for two HRTs are given in (Table 5). As can be seen, the HRT of 4.5d gave higher COD solubilization and released more ammonia and phosphate than the HRT 3d. Moreover, the VFA yield per gram of volatile solids, 0.57, for HRT 4.5d was significantly higher than 0.5 for HRT 3d (t=-2.94, df=8, ***P***=.02, CI=95%).

**Table 5.** Performance parameters of two different operational conditions used in mesophilic acidogenic fermentation.

| HRT (Days) | Solubilization ($\Delta g$-<br>SCOD/g-$VS_0$) ±<br>SD | $Y_{VFA}$ ($\Delta g$-<br>VFA/g-$VS_0$) ±<br>SD | Ammonia<br>Release (%) ±<br>SD | Phosphate<br>Release<br>(%) ± SD |
| --- | --- | --- | --- | --- |
| 4.5<sup>a</sup> | 0.28 ± 0.06 | 0.57 ± 0.06 | 35 ± 10.74 | 13.7 ± 8.77 |
| 3<sup>b</sup> | 0.19 ± 0.05 | 0.50 ± 0.06 | 29 ± 0.11 | 11 ± 0.06 |

a: 9 measurements for solubilization, $Y_{VFA}$ , 8 measurements for ammonia and phosphate release, b: 9 measurements for solubilization, $Y_{VFA}$ , 7 measurements for ammonia and phosphate release

In the biopolymer synthesizing process, the aim was to generate a stable VFA weight ratio distribution with a high VFA/SCOD ratio for an efficient PHA synthesis during the whole process. Concisely, the VFA stream with a higher dominance of even numbers of carbon atom acids means a higher 3-hydroxybutyrate (HB) monomer synthesis compared to the 3-hydroxyvalerate (HV), which is correlated with the net prevalence of odd numbers of carbon atom acids (propionic, valeric, and isovaleric acid) [32]. As can be inferred, the stability in the VFA spectrum means a predictable and reproducible PHA monomer production. Accordingly, the physical and mechanical features of synthesized biopolymers are stable [33,34].

(Figure 5) reports the weight ratio distribution of the VFAs for two HRTs. The main fractions were acetic acid (38-42%), butyric acid (24%), caproic acid (16-18.5%), propionic acid (9-11%), and valeric acid (5%). This VFA distribution, with a major part of butyric and acetic acid, is in line with those reported in similar studies [28,30]. In this respect, VWRD is determined by the type of feedstock and food waste rather than the operational conditions.

**Figure 5.**
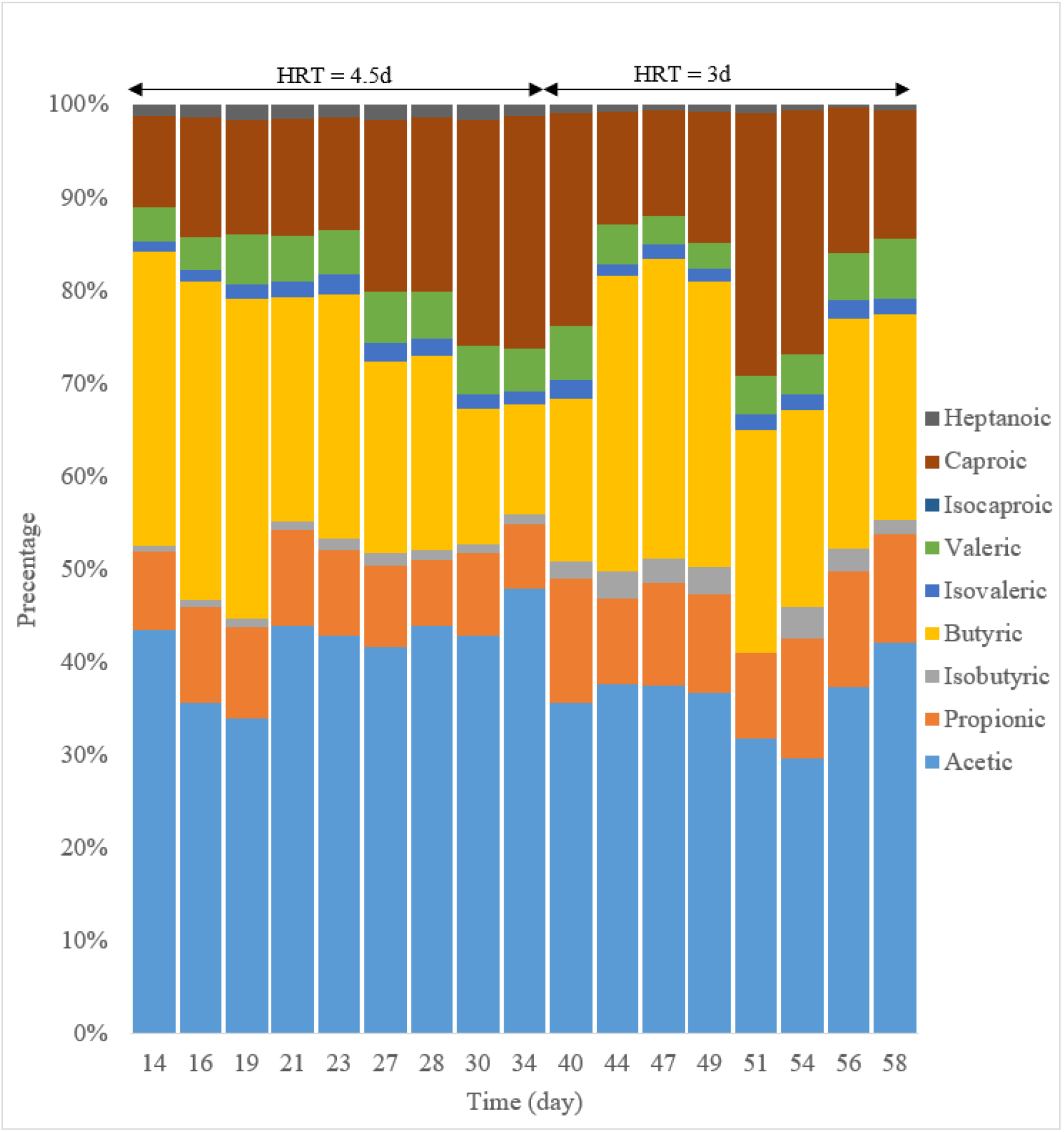
VFAs weight ratio distribution for mesophilic acidogenic fermentation

### Anaerobic digestion

(Table 6) summarizes the performance parameters and the results from the kinetics study for anaerobic digestion. This study obtained a remarkably high value for hydrolysis rate (i.e., 0.58 1/d) with no lag phase. Besides, the biogas yield of 0.61-0.83 (g-biogas/g-VS), the SMP of 0.133-0.204-), and an average composition of 45%-58% methane (v-/v-biogas) were obtained. According to (Figure 6), adding biochar provided the desirable conditions for the growth of hydrogen using methanogenesis manifested through a higher maximum volumetric methane content (86% vs 66%, v/v).

**Table 6.** The performance indicators for anaerobic digestion and results from the kinetics study for two models: (1) First-Order rate and (2) Modified Gompertz.

| Experiments | SMP<sup>a</sup> ($\text{CH}_4\text{-Nm}^3/\text{kg} - \text{VS}$) | SGP<sup>b</sup> ($\text{CH}_4\text{-Nm}^3/\text{kg} - \text{VS}$)±SD | K<sup>c</sup> (1/d) | R<sub>m</sub><sup>d</sup> ($\text{CH}_4\text{-ml/g-VS} \cdot \text{d}$) |
| --- | --- | --- | --- | --- |
| Without biochar | 0.204 | 0.540 | 0.57 | 76.12 |
| Biochar (0.12 g-biochar/g-VS<sub>added</sub>) | 0.133 | 0.567 | 0.69 | 62.42 |

| Experiments | $\lambda^e$ (d) | RMSE<sup>f</sup> FO<br>($CH_4$-<br>$Nm^3/kg - VS$) | RMSE MG<br>($CH_4$-<br>$Nm^3/kg - VS$) | Max CH4 content<br>(%, v/v) |
| --- | --- | --- | --- | --- |
| Without biochar |  |  |  |  |
|  | 0 | 10.4 | 6.82 | 68.5 |
| Biochar (0.12 g-biochar/g-<br>$VS_{added}$) | | | | |
|  | 0 | 5.74 | 5.59 | 86 |
| Biochar (0.24 g-biochar/g-<br>$VS_{added}$) | | | | |
|  | 0 | 9.64 | 3.39 | 76.5 |

a: specific methane production, b: specific gas production, c: hydrolysis rate, d: maximum methane production rate, e: lag phase, and f: root mean squared error

**Figure 6.**
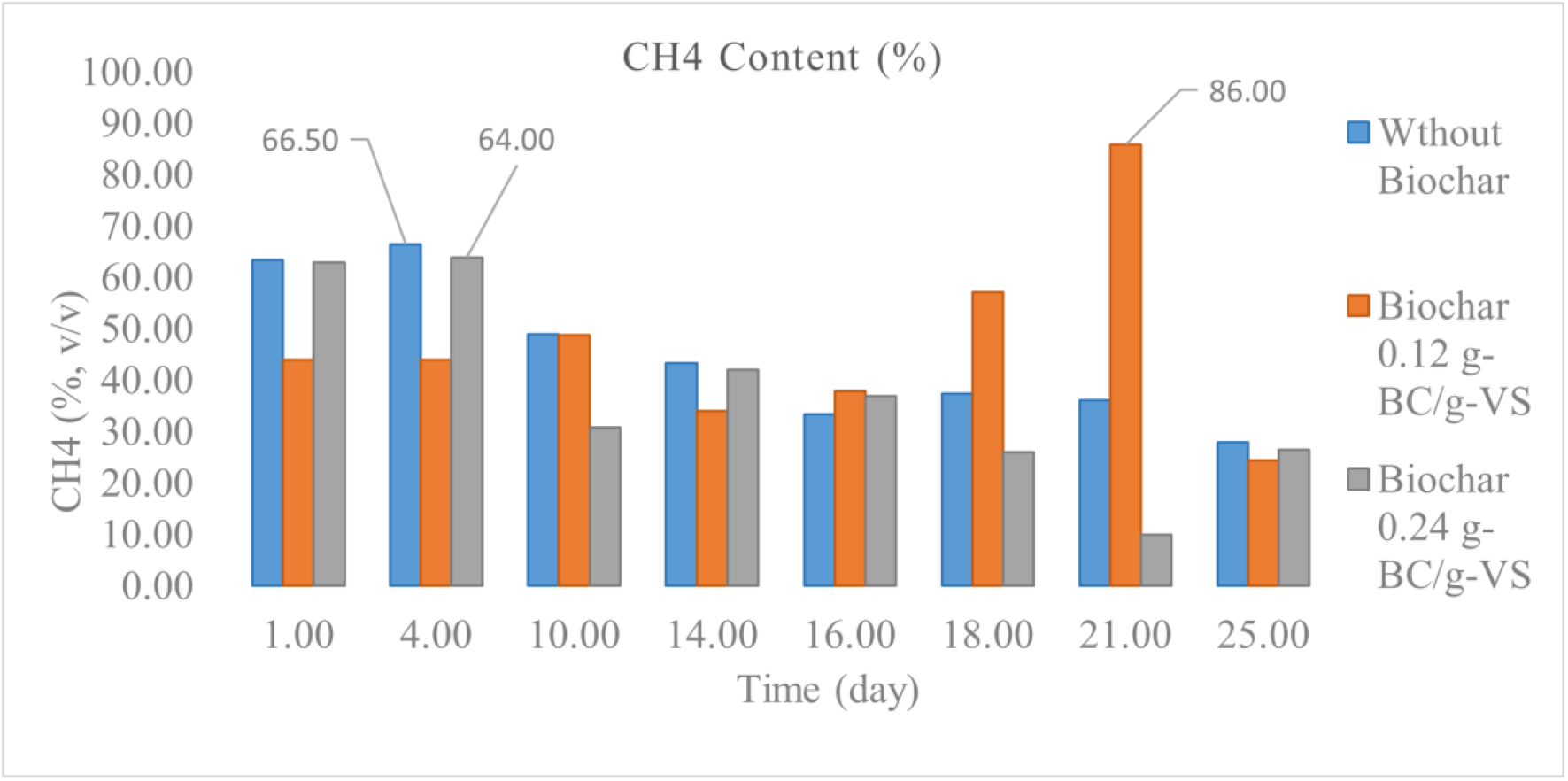
content in volume basis (v/v) for 3 different biochar dosages in anaerobic digestion

The mass flow model was adopted for 0.12 (g-biochar/g-VS) as the only feasible solution. Unlike other dosages, it could satisfy the assumptions for FS/IN 0.3 at HRT 20d, which was adequately long enough to let the methanogens reproduce themselves. Detailed information is available in an EXCEL sheet named “DIGESTER DESIGN”. Besides, the high alkalinity of the biochar as reported in the supplementary documents (Table 1) signifies a benefit of the biochar addition in limiting the concern over pH drop for high OLR in full-scale implementation. Accordingly, almost four-fold the ordinary OLR was obtained, i.e., 6.25 (kg-VS/, by a minimum water dilution, knowing that the biochar could maintain the stability of the process. Therefore, the digester volume will decline at the rate of 28 L/PE. Hence, the presented mass flow line model was implemented based on the results from 0.12 (g-biochar/g-VS), the weighted average composition of biomethane as 35% (v/v), and the SGP as 0.56 (biogas-) for HRT 20d corresponding to FS/IN of 0.3.

Based on the root mean squared error (RMSE) reported in (Table 6), both models were almost identical for describing biomethane production for biochar dosage of 0.12 (g-biochar/g-VS), and for the simplicity we used the FO rate model in the feasibility study.

### Technical and economic assessment

Assuming an imaginary municipality of 70,000 (PE) and the amount of TS production per capita as 0.3 (kg/PE) per day [35], the inlet to the scale-up line would be 21,000 (kg-TS/d).

In the first scenario, the biowaste stream, after passing through the screw-press and pretreatment unit, had a mass flow of 113,788 (kg/d), TS of 4.1% (kg/kg), and VS of 3.1% (kg/kg). Then, the mixture was heated to 37° before and in the acidogenic fermenter, which was operated at HRT = 4.5d and OLR of 6.89 (kg-VS/m^3^.d). This process was performed to convert biosolids into the VFAs and SCOD at the concentration level of 30.77 (g-SCOD/L) and 34 (g-SCOD/L), respectively. At this step, the gaseous flow rate was assumed to be zero as HRT of 4.5d is quite short for any adequate growth of methanogens in mesophilic conditions. The stream out of the acidogenic fermenter had a mass flow rate of 113,788 (kg/d), with a VFA content of 3501 (kg-SCOD/d), which could be used in the PHA synthesizing step [36]. The outlet of this step was used in the separator to gain overflow and the solid cake. Later, the solid cake was minimally diluted by water before being fed into a mesophilic anaerobic digester with a biochar addition of 0.12 (g-biochar/g-VS). The anaerobic digester received a TS content of 18% (kg/kg) and a flow rate of 18,180 (kg/d) corresponding to HRT 20d and OLR of 6.25 (kg-VS/m^3^.d). Overall, an SGP of 0.285 (Nm^3^ /kg-VS) was obtained assuming zero gas production in acidogenic fermentation.

In the second scenario, the fresh feedstock, after being screw-pressed, had a mass flow rate of 4,678 (kg-TS/d) and 28% (kg/kg) dry matter. Then, it was diluted with water and heated before being fed into the anaerobic digester. At this step, the mass flow rate of 85,012 (kg/d) with a TS of 6% (kg/kg) entered the digester with a volume of 2125,(m^3^) leading to an HRT of 25d and OLR of 1.7 (kg-VS/m^3^.d). The SMP of 0.311 (Nm^3^ -biogas /kg-VS) was obtained by destroying 80% of the VS.

In this study, working volumes of 512 (m^3^) and 364 (m^3^)were adopted for the acidogenic fermenter and anaerobic digester in the first scenario, respectively, and 2,125 (m^3^)for the anaerobic digester in the second scenario. As a result, the capital cost for the presented line was almost €809,000, roughly half of the quantity for the single-step anaerobic digestion (Figure 7). Unlike the single-step anaerobic digestion that converts all VS to biogas, this novel line shared the recovery of VS between a higher added-value VFAs and biogas production and expectedly generated 10-fold higher benefits, €375,085. Consequently, the payback period was reduced by more than 20 times in 2 years (Figure 7). This period was achieved at the cost of less amount of surplus energy, 2251 MJ/d for two-step fermentation vs. 21567 MJ/d for single-step anaerobic digestion (Figure 8).

**Figure 7.**
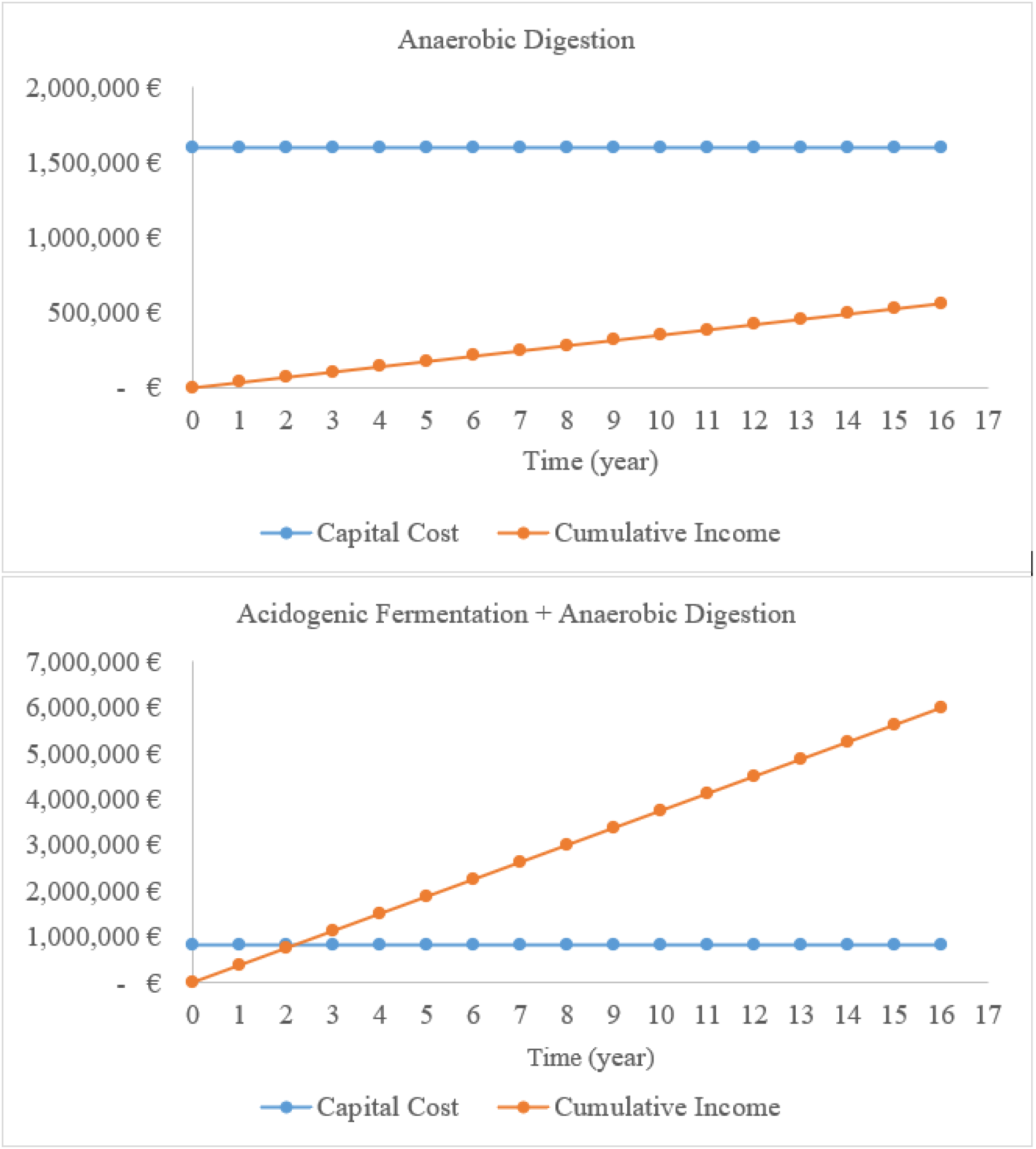
Capital cost and cumulative yearly income for two proposed scenarios

**Figure 8.**
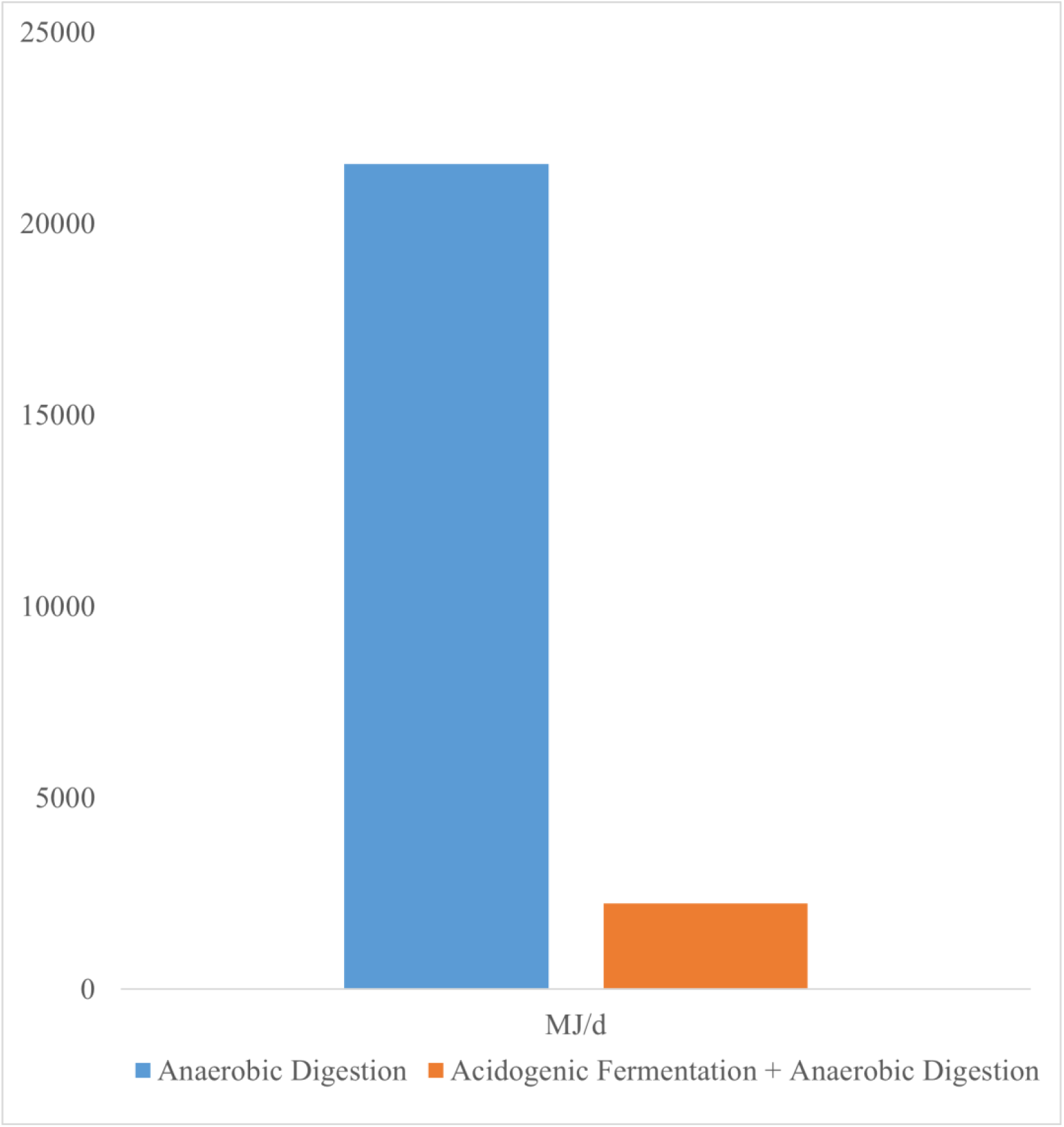
Daily net energy production for two scenarios

## Discussion

### Principal Results

We showed that multistep fermentation is both economically and technically feasible. The findings indicated that producing VFAs and biogas in separate stages can significantly reduce the payback period for upcoming investments in biorefinery projects and result in the creation of a highly desired stream that is rich in VFAs. Additionally, the process stability could be maintained even in high OLR by adding biochar and converting the VS’s easily biodegradable COD content into VFAs in the first phase. This would preserve energy and water and reduce the digester’s volume.

### Comparison with previous studies

Because of the extra pretreatment unit in the present research, the VFA yield of 0.57-0.63 (Δg-VFA/g-) was roughly double the value reported by Valentino et al. [30] for the same OMSW.

Our results also indicate a spectacular improvement in the process kinetics, which was manifested through more than an 8-fold rise in the hydrolysis rate (0.58 vs. 0.07, 1/d) and a full decrease in lag phase (0 vs. 2.69, d) as opposed to the previous study by Karki et al. [37]. This improvement is attributed to the destruction of the solids structure caused by bacterial enzymes and a hot alkaline solution. Additionally, a higher active biomass per feedstock was provided using a fine-tuned FS/IN ratio of 0.3 (VS basis), which was noticeably lower than the quantities (1 and 0.5) reported in similar studies [37,38].

The values for SMP and mean methane volumetric content presented in this study are lower than those reported by Valentino et al. (28), i.e., 0.25 (CH_4_-Nm^3^/Kg-VS) and 63%-64% (v/v). This difference is explained by the added fresh WS, which has a higher digestible content and better nutrient balance than the fermented solids. Similarly, the SMP in this study was lower than 0.384(CH_4_-Nm^3^/Kg-VS) in the study by Moreno et al. [38]. This author investigated the anaerobic digestion of residual solids from two steps of bioethanol production and saccharification on OMSW. In this respect, bioethanol production can convert only part of cellulose, starch, and some dissolved carbohydrates. Consequently, a great part of the biosolids’ volatile content, nearly 70%, is still available to be harvested in different biorefinery schemes compared with the one proposed in this method with 34%. Besides, the fermented OMSW would have a completely incompatible composition since it did not only come from different geographical locations (Spain and the UK) with different food habits but also underwent different biological pretreatment.

The multistep recovery line proposed in this study was more technically practicable than the sequential conversion of OMSW to bioethanol and biogas proposed in the study by Moreno et al. [38]. As it requires sterilization conditions, imposing an additional operational cost and the bioethanol concentration should be high enough to lower the cost of the subsequent distillation step.

Furthermore, our method for VFA production distinctively from biogas was preferable to the study by Papa et al. [9], wherein the operational alteration on a single anaerobic digester was performed to obtain VFAs and biogas. These researchers asserted that the single-step recovery of biogas and VFAs was feasible by increasing OLR while keeping the SMP of the reactor almost unaffected. The main recovery path for the VS was still biogas production in their study, which accounted for more than 90% of the VS conversion. Meanwhile, the present study obtained 47% and 32% of the biogas and VFA conversion share, respectively. Further, whereas the destruction of VS of around 70% was achieved in both studies, their proposal limited the VFA distribution to propionic and butyric acid. The explanation is that some of the VFAs were converted into biogas in the same unit, which could negatively affect the PHA synthesis step later.

### Conclusion and limitations

This paper demonstrated the technical and economic feasibility of multistep fermentation. The results of this study indicate that the production of VFAs and biogas in distinct steps can considerably shorten the payback period for future investments in biorefinery projects and produce a highly desirable VFA-rich stream. Further, adding biochar and converting easily degradable COD content in the VS into VFAs in the first step could maintain the process stability even in high OLR. As a result, it leads to energy and water preservation and a decrease in the digester volume. However, consideration should be paid to the full-scale implementation since the pilot studies cannot resemble the stability of the real process. For instance, operational alterations such as raising OLR and the addition of biochar in the full-scale implementation might perturb the process of pH or the synergetic balance between the bacterial communities and stop the process completely, which were never observed in our experimental study. Further, the superb profitability of the proposed line was highly variable because our cost analysis was too simplistic and did not elaborate on all the possible associated expenditures. Besides, since many of its components were from subject matter experts rather than the pilot studies budget, they were prone to site variations and uncertainties. Addressing the systematic uncertainty in the labor and material costs due to the changes in the supply chain issues, inflation, and site variations is beyond our scope. Moreover, caution should also be considered regarding the significance of the BMP results with the marginal difference since the number of samples was not large enough for statistical analysis. Nevertheless, the results presented in this study were prepared technically and financially cautiously to encourage the revolution in the current state of organic waste valorization in Italy and any similar location.

In conclusion, a robust framework was proposed to assess the valorization of organic waste through experimental tests, statistical analysis, process kinetics, and mass and energy flow analysis. The findings support considerably higher profitability and, thus, a shorter payback period for the multistep fermentation than the current single anaerobic digestion. Additionally, our results encourage the circular economy perspective on converting OMSW into biogas and VFAs with the pros of fewer residual solids due to reusing them in a pyrolysis line.

## Supporting information

Biomethanation Potential Test, Mass Flow Analysis, Cost Analysis, Anaerobic Digester Design, Laboratory Data, Satistical Input Data, Kinetics Study

Tables, Equations, and Codes

## Acknowledgments

The authors gratefully acknowledge Francesco Valentino for providing helpful comments and guidance during the study. We also appreciate Alessio Dell ′Olivo and Aditi Parmar Chitharanjan for helping with laboratory experiments and Material flow analysis. We also thank Marco Gottardo for his helpful input on gas chromatography.

The authors gratefully acknowledge the Italian Ministry of University and Research (MUR) and the University of Ca′ Foscari for financially supporting this research in the Frame of Programma Operativo Nazionale Ricerca e Innovazione (PON).

## Multimedia Appendix 1

## Abbreviations and Nomenclature

mg: Milligram
g: Gram
kg: Kilogram
µm: micrometer
m: Meter, µ
L: microliter
ml: milliliter
L: Liter
Nm^3^: Normal Cubic Meter
MJ: Mega Joule
v/v: Volumetric Basis
CH_4_: Methane
N: Nitrogen
P: Phosphorous
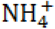: Ammonium
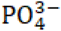: Phosphate
PE: Person Equivalents
FS: Feedstock
IN: Inoculum
BMP: Bio-methanation Potential Test
CI: Confidence Interval
COD: Chemical Oxygen Demand
EU: European Union
FO: First-Order
HB: 3-hydroxybutyrate
HRT: Hydraulic Retention Time
HV: 3-hydroxyvalerate
LOD: Limit of Detection
MG: Modified Gompertz
N: Total Nitrogen
OLR: Organic Loading Rate
OMSW: Organic Municipal Solid Waste
PHA: Polyhydroxyalkanoates
P: Total Phosphor
RMSE: Root Means Squared Error
rpm: round per minute
SCOD: Soluble Chemical Oxygen Demand
SGP: Specific Gas Production
SMP: Specific Methane Production
TS: Total Solids
TKN: Total Kjeldahl Nitrogen
VFA: Volatile Fatty Acid
VS: Volatile Solids
VWRD: VFA Weight Ratio Distribution:
WWTP: Wastewater Treatment Plant
WS: Waste Sludge
K: Hydrolysis rate
R_m_: Maximum Methane Production Rate
λ: Lag Phase
N_2_: Nitrogen
O_2_: Oxygen
CO:_2_: Carbon dioxide
NaOH: Sodium Hydroxide

