## Supplementary material for "Converting Organic Municipal Solid Waste into Volatile Fatty Acids and Biogas: Experimental Pilot and Batch Studies with Statistical Analysis": Tables, Equations, and Codes

### Carbonaceous Characteristics

(Table 1) reports the values for physical and chemical characteristics of the biochar in our study.

Table 1. Chemical and physical characteristics of biochar

| **Biochar Characteristics** | **Weight Ratio (g/kg)** | **Weight Percentage (%)** |  |  | **pH** |
| --- | --- | --- | --- | --- | --- |
| Total Nitrogen |  |  |  |  |  |
|  | **-** | <0.5% |  |  | **-** |
| 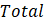Potassium |  |  |  |  |  |
|  | 3.020 | **-** |  |  | **-** |
| Total Phosphor |  |  |  |  |  |
|  | - | 0.03% |  |  | **-** |
| Total Calcium |  |  |  |  |  |
|  | 9.920 | **-** |  |  | **-** |
| Total Magnesium |  |  |  |  |  |
|  | 0.852 | **-** |  |  | **-** |
| Total Sodium |  |  |  |  |  |
|  | 0.291 | **-** |  |  | **-** |
| Total carbon of biological origin on dry matter |  |  |  |  |  |
|  | - | 65% |  |  | **-** |
| pH |  |  |  |  |  |
|  | - | **-** |  |  | 9.85 |
| Ash content |  |  |  |  |  |
|  | - | 5.50% |  |  | **-** |
| Carbonia from carbonates |  |  |  |  |  |
|  | - | <0.1% |  |  | **-** |

### Equations for performance parameters and kinetics Study

The COD solubilization, VFA yield, and VFA rate were calculated based on their ultimate concentrations in (g/L), and the initial values for VFA and SCOD, represented by
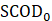
(g/L) and
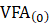
 (g/L), plus the reported values for initial volatile solds content,
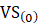
 (g/g). Similarly, the release for ammonia and phosphate were determined based on their final concentrations (g/L) and the nitrogen and phosphor content available in the unfermented total solids part, reported for TKN (g/kg), P (g/kg), and
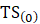
 (g/kg). In following, you can see the equations used in this study:

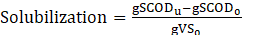

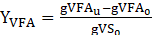

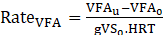

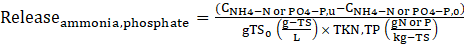

The equations for First-Order rate and Modified gompertz model is outlined as following:

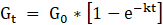
 first-order

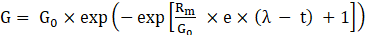
 modified Gompertz model

Where
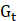
 is cumulative methane yield at time t (mL/g-
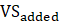
), K: methane production rate constant (first order disintegration rate constant) (1/day); t: Time (days). Moreover, in the modified Gompertz model,
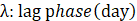
,
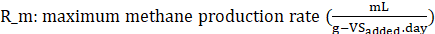
.

### Digester Design

We search for specific HRT and Q to reach the FS/IN ratio of 0.3 and meeting the following criteria:

1. Avoid an extremely high OLR >15 (kg.VS/m3.d) to maintain the stability of the process with respect to pH

2. No Washing out of anaerobes bacteria could happen HRT>12 days

3. keep the FS/IN ratio close to 0.3

We did calculation as follow:

HRT =
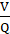

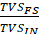
 =
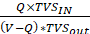

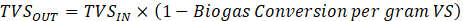

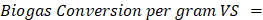

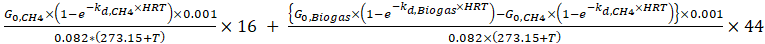

### KNN Imputation Code

We wrote the following code to fill the values for gas contents on days when there was no measurements conducted.

\# This code is used for prediction of biomathane content in the days

\# where data are not measured using K-Nearest Neighbour Hood Estimation

from sklearn.impute import SimpleImputer

import numpy as np

from sklearn.impute import KNNImputer

import matplotlib.pyplot as plt

#bottle 1 and 2

data = [[0, 0], [1, np.nan], [2, np.nan], [3,np.nan], [4, np.nan],[6, 56], [7, np.nan], [8,np.nan], [10,44.65], [11,np.nan], [14,43.93], [16,57.5], [18,40], [21,48], [23,np.nan], [25,56.28]]

data_train = [[0, 0], [6, 56], [10,44.65],[14,43.93],[16,57.5], [18,40], [21,48], [25,56.28]]

data_test = [[1, np.nan], [2, np.nan], [3,np.nan], [4, np.nan],[7, np.nan], [8,np.nan], [11,np.nan],[23,np.nan]]

print("\n")

print("====== Data for Bottle 1 and 2 - Only Inoculum ======\n")

print('Data =',data)

imputer = KNNImputer(n_neighbors=4, weights='distance')

imputer.fit(data_train)

imputer_KNN=imputer.transform(data_test)

print(imputer_KNN)

#bottle 3 and 4

data = [[0, 0], [1, 63.5], [2, np.nan], [3,np.nan], [4, 66.5],[6, 46], [7, np.nan], [8,np.nan], [10,49], [11,np.nan], [14,43.4], [16,33.5], [18,37.5], [21,36.1], [23,np.nan], [25,28]]

data_train = [[0, 0],[1, 63.5],[4, 66.5],[6, 46],[10,49],[14,43.4],[16,33.5],[18,37.5], [21,36.1], [25,28]]

data_test = [[2, np.nan], [3,np.nan], [3, np.nan],[7, np.nan], [8,np.nan],[11,np.nan], [23,np.nan] ]

print("\n")

print("====== Data for Bottle 3 and 4 Without Biochar + Inoculum + FS ======\n")

print('Data =',data)

imputer = KNNImputer(n_neighbors=4, weights='distance')

imputer.fit(data_train)

imputer_KNN=imputer.transform(data_test)

print(imputer_KNN)

#bottle 5 and 6

data = [[0, 0], [1, 44], [2, np.nan], [3,np.nan], [4, 44],[6, 44.55], [7, np.nan], [8,np.nan], [10,48.9], [11,np.nan], [14,34], [16,38], [18,57.2], [21,86], [23,np.nan], [25,24.5]]

data_train = [[0, 0], [1, 44], [4, 44], [6, 44.55],[10,48.9],[14,34],[16,38],[18,57.2], [21,86], [25,24.5]]

data_test = [[2, np.nan], [3,np.nan], [7, np.nan], [8,np.nan],[11,np.nan], [23,np.nan] ]

print("\n")

print("====== Data for bottle 5 and 6 With Biochar 0.12 g-BC/g-VS + Inoculum + FS ======\n")

print('Data =',data)

imputer = KNNImputer(n_neighbors=4, weights='distance')

imputer.fit(data_train)

imputer_KNN=imputer.transform(data_test)

print(imputer_KNN)

#bottle 7 and 8

data = [[0, 0], [1, 63], [2, np.nan], [3,np.nan], [4, 64],[6, 42.5], [7, np.nan], [8,np.nan], [10,30.8], [11,np.nan], [14,42.15], [16,37], [18,26], [21,10], [23,np.nan], [25,26.5]]

data_train = [[0, 0], [1, 63],[4, 64],[6, 42.5],[10,30.8],[14,42.15],[16,37],[18,26], [21,10], [25,26.5]]

data_test = [[0, np.nan], [2, np.nan], [3,np.nan], [4, np.nan],[7, np.nan], [8,np.nan],[11,np.nan], [23,np.nan] ]

print("\n")

print("====== Data for bottle 7 and 8 with Biochar 0.24 g-BC/g-VS + Inoculum + FS ======\n")

print('Data =',data)

imputer = KNNImputer(n_neighbors=4, weights='distance')

imputer.fit(data_train)

imputer_KNN=imputer.transform(data_test)

print(imputer_KNN)

### Statistical Analysis code

#This project was done to visualize the

#the range of variation of the VFA/sCOD and VFA concentration and Yield

#for two HRTs of 4.5 and 3 days. It also

#applies the statistical analysis to the data assuming normality

\# and homogenity of variance along with pairedness between the data

#Read files of the data

VFA = read.table("D:\\Academy\\Articles\\Biogas and VFA\\Mesophilic My data\\JMIRX\\Supplementary Info\\data of mesophilic test-HRT4.5,3.csv",dec = ".",sep = ",",header=TRUE)

VFA=VFA[,!names(VFA) %in% c("X", "X.1",'X.2')]

VFA<-na.omit(VFA)

VFA

head(VFA)

myColors <- c(rgb(0.1,0.1,0.7,0.5) ,

rgb(0.8,0.1,0.3,0.6))

attach(VFA)

boxplot(HRT.3d,HRT.4.5d, xlab='HRT',ylab='VFA/SCOD',col = myColors, names=c('HRT3d', 'HRT4.5d'))

boxplot(VFA4.5d, VFA3.d, xlab='HRT', ylab= 'VFA(mgSCOD/l)', col=myColors, names=c('HRT3d', 'HRT4.5d'))

boxplot(Y4.5d, Y3d, xlab='HRT', ylab= 'Yield VFA(g-VFA/g-VS)', col=myColors, names=c('HRT3d', 'HRT4.5d'))

\# Visualization of Histogram for two given HRTs

hist(HRT.3d)

hist(HRT.4.5d)

hist(VFA3.d)

hist(VFA4.5d)

hist(Y4.5d)

hist(Y3d)

\# Printing mean and standard deviation of VFA/SCOD HRT=4.5d

mean(HRT.4.5d)

sd(HRT.4.5d)

\# Printing mean and standard deviation of VFA/SCOD HRT=3d

mean(HRT.3d)

sd(HRT.3d)

\# Printing mean and standard deviation of VFA HRT=4.5d

mean (VFA4.5d)

sd(VFA4.5d)

\# Printing mean and standard deviation of VFA HRT=4.5d

mean (VFA3.d)

sd(VFA3.d)

\# Printing mean and standard deviation of Y_VFA HRT=4.5d

mean (Y4.5d)

sd(Y4.5d)

\# Printing mean and standard deviation of Y_VFA HRT=3d

mean (Y3d)

sd(Y3d)

#t-test for VFA/SCOD production

t.test(x=HRT.3d, y=HRT.4.5d, paired = TRUE, conf.level = 0.95)

#t-test for VFA concentration

t.test(x=VFA3.d, y=VFA4.5d, paired = TRUE, conf.level = 0.95)

#t-test for Y_VFA concentration

t.test(x=Y3d, y=Y4.5d, paired = TRUE, conf.level = 0.95)
